# Investigation on the requirements for YbbN/CnoX displaying thiol-disulfide oxidoreductase and chaperone activities

**DOI:** 10.1101/2020.04.09.034579

**Authors:** Diogo de Abreu Meireles, César Henrique Yokomizo, Luís Eduardo Soares Netto

## Abstract

YbbN/CnoX are proteins that display a Trx domain linked to a tetratricopeptide (TPR) domain, which are involved in protein-protein interactions and protein folding processes. YbbN from *Escherichia coli* (*Ec*YbbN) displays a co-chaperone (holdase) activity that is induced by HOCl (bleach). *Ec*YbbN contains a SQHC motif within the Trx domain and displays no thiol-disulfide oxidoreductase activity. *Ec*YbbN also presents a second Cys residue at Trx domain (Cys63) 24 residues away from SQHF motif that can form mixed disulfides with substrates. Here, we compared *Ec*YbbN with two other YbbN proteins: from *Xylella fastidiosa* (*Xf*YbbN) and from *Pseudomonas aeruginosa* (*Pa*YbbN). While *Ec*YbbN displays two Cys residues along a SQHC[N_24_]C motif; *Xf*YbbN and *Pa*YbbN present two and three Cys residues in the CAPC[N_24_]V and CAPC[N_24_]C motifs, respectively. These three proteins are representatives of evolutionary conserved YbbN subfamilies. In contrast to *Ec*YbbN, both *Xf*YbbN and *Pa*YbbN: (1) reduced an artificial disulfide (5,5′-dithiobis-(2-nitrobenzoic acid) = DTNB); and (2) supported the peroxidase activity of Peroxiredoxin Q from *X. fastidiosa*, suggesting that *in vivo* these proteins might function similarly to the canonical Trx enzymes. Indeed, *Xf*YbbN was reduced by *Xf*Trx reductase with a high catalytic efficiency (*k_cat_*/K_m_=1.27 × 10^7^ M^−1^.s^−1^), like the canonical *Xf*Trx (*Xf*TsnC). Furthermore, *Ec*YbbN (as described before) and *Xf*YbbN, but not *Pa*YbbN displayed HOCl-induced holdase activity. Remarkably, *Ec*YbbN gained disulfide reductase activity while lost the HOCl-activated chaperone function when the SQHC was replaced by CQHC. In contrast, the *Xf*YbbN C40A mutant lost the disulfide reductase activity, while kept its HOCl-induced chaperone function. Finally, we generated a *P. aeruginosa* strain with the *ybbN* gene deleted, which did not present increased sensitivity to heat shock or to oxidants or to reductants. Altogether, our results suggest that different YbbN/CnoX proteins display distinct properties and activities, depending on the presence of the three conserved Cys residues.

**Graphical Abstract:** 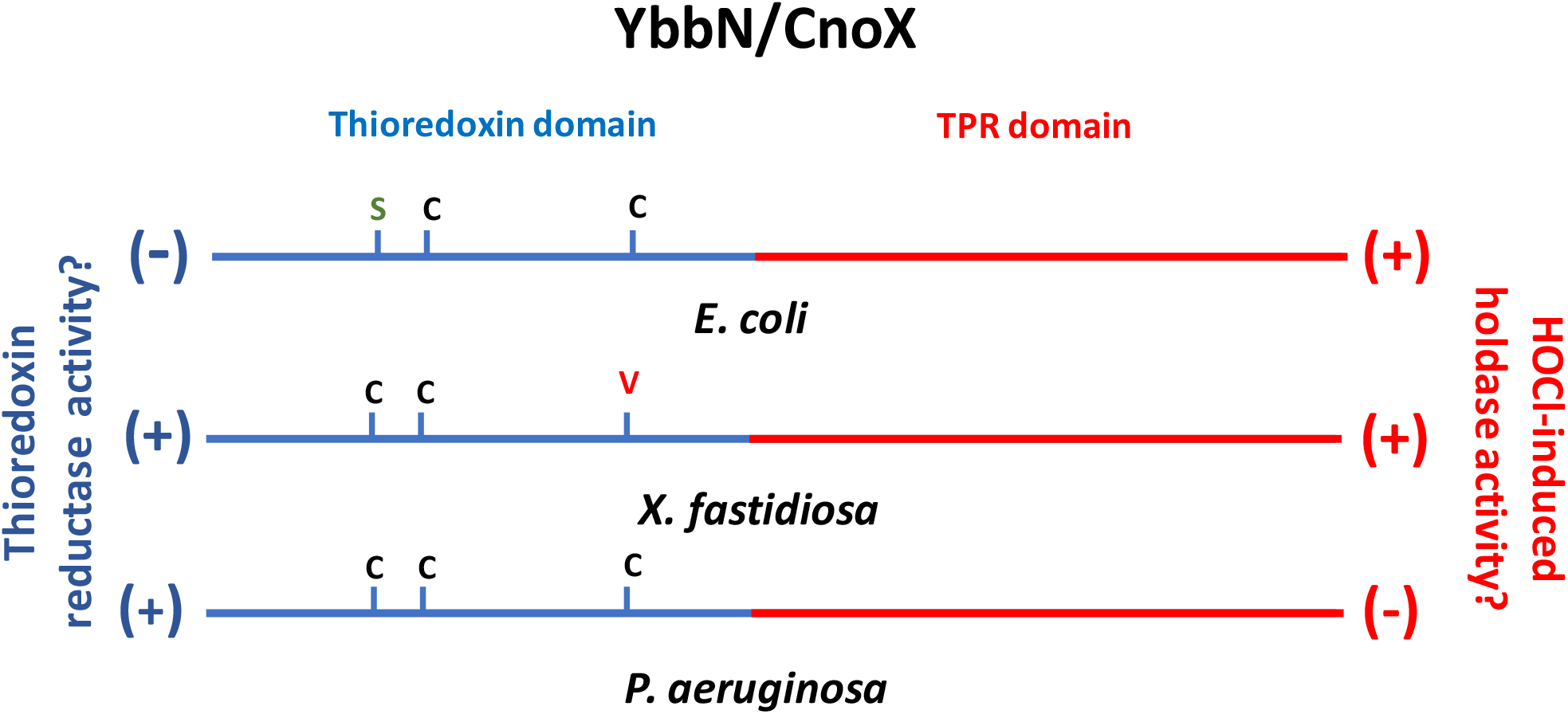

**Highlights:**

- CXXC motif is required for the thiol-disulfide reductase activity of YbbN proteins.
- *Xf*YbbN and *Pa*YbbN display thiol-disulfide oxidoreductase activity
- The affinities of *X*fTrxR for *Xf*YbbN and *Xf*TsnC (canonical Trx) are comparable
- *Xf*YbbN and *Ec*YbbN, but not *Pa*YbbN, display holdase activity induced by hypochlorous acid
- Engineering *Ec*YbbN/CnoX by inserting a Cys residue in the SQHC motif resulted in a gain of function (thiol-disulfide oxidoreductase activity) and abolished the HOCl-induced holdase activity.

## 1. Introduction

Thioredoxin (Trx) enzymes are widely distributed among organisms, presenting key roles in cells such as deoxyribonucleotide synthesis and as substrates for enzymes such as peroxiredoxins, methionine sulfoxide reductase and phosphoadenosine phosphosulfate reductase (PAPS) [1]. In addition, Trx also participates in redox signaling pathways [2,3]. Trx enzymes are also present fused to other domains, but their cellular roles are still poorly studied. The diversity of the domains fused with Trx-like proteins is very high, with more than 300 distinct partners already identified [4]. YbbN proteins are one of these multi-domain proteins that contain a Trx domain at the N-terminal portion (12 KDa) and a tetratricopeptide repeat (TPR) domain at the C-terminal portion (20 KDa). TPR domain is also widely distributed, composed from 3 to 16 repeats, each one containing around 34 amino acids [5]. This domain provides a concave groove for protein-protein interactions [6,7]. Noteworthy, several proteins containing the TPR domain are involved in virulence [8]. So far, among all the YbbN enzymes, only *Ec*YbbN (PDB Id = 3QOU) [6] and YbbN from *S. typhimurium* (PDB Id = 3QDN), both presenting SQHC motifs, had their structures elucidated.

The Trx domain of *Ec*YbbN lacks the most N-terminal residue of the typical CXXC motif (SQHC) and as expected this protein displays no oxidoreductase activity [9]. *Ec*YbbN is involved in the regulation of chaperone machinery, possibly by protein-protein physical interactions through its TPR motifs and with the assistance of GroEL [5, 6]. Indeed, *E. coli ybbN*-deficient strain (Δ*ybbN*) is sensitive to heat stress but not to H_2_O_2_ stress [10]. More recently, *Ec*YbbN was described as a key protein in bacterial response to hypochlorous acid (HOCl) [11]. HOCl induces the chlorination of specific amino acid residues at TPR domain, activating a holdase activity. Furthermore, *Ec*YbbN displays a second activity based on Cys63, which is 24 residues away from the SQHC motif, still in the Trx domain. *Ec*YbbN forms mixed disulfides with target proteins through its Cys63 residue and glutathione (GSH) is required to resolve these *Ec*YbbN-substrate complexes [11]. Therefore, *Ec*YbbN presents dual function: preventing irreversible protein aggregation and protection of cellular proteins from hyperoxidation.

In this work, we characterized enzymes from *E. coli*, from *Xylella fastidiosa* and from *P. aeruginosa*. *X. fastidiosa* and *P. aeruginosa* are two gram-negative bacteria, the former is a plant pathogen that produces a significant economic impact in the citri, grape and olive cultures [12] and the latter is an important opportunistic human pathogen [13,14]. The expression of YbbN from *X. fastidiosa* (*Xf*YbbN) and from *P. aeruginosa* (*Pa*YbbN) are both induced by high temperatures [15,16], which suggest that *Xf*YbbN and *Pa*YbbN might function as co-chaperones. There are no studies on the putative oxidoreductase activities of *Xf*YbbN and *Pa*YbbN.

In this work, we initially analyzed the distribution of YbbN proteins as a function of the presence of Cys residues in a CXXC motif and considering the amino acid present in the position equivalent to Cys63 of *Ec*YbbN. Accordingly, we propose here that YbbN proteins should be divided in four functional groups. Then, we compared the activities of *Xf*YbbN and *Pa*YbbN that contain the canonical WCXPC motif with *Ec*YbbN. Furthermore, we describe that the deletion of *ybbN* gene in *P. aeruginosa* did not render cells more sensitive to HOCl. Our data indicated that the presence or absence of Cys residues in the CXXC motif or 24 residues far away from this redox center are relevant factors that modulate the three activities reported for YbbN enzymes: (1) co-chaperone/holdase; (2) thiol-disulfide oxidoreductase; (3) prevention of hyperoxidation by formation of mixed disulfides.

## 2. Materials and methods

### 2.1 Strains and growth conditions

*P. aeruginosa* UCBPP-PA14 [13] and *E. coli* BW25113 [17] were used as wild type strains, except when otherwise indicated. These bacteria were grown in Lysogenic Broth (LB) medium at 37°C supplemented with appropriated antibiotics: carbenicillin (300 μg/mL), ampicillin (100 μg/mL) or kanamycin (15 μg/mL). The plasmids and strains used in this work are listed in Supplementary Table 1.

The *ybbN* deletion strain of *E. coli* BW25113 was acquired from Keio collection mutants [17]. The kanamycin resistance cassette was excised using FLP-FRT recombination system and the colonies that had lost kanamycin resistance were checked by PCR using oligonucleotides described in the Supplementary Table 1. A clean, *in-frame ybbN* deletion strain of *P. aeruginosa* was obtained through homologous recombination. Briefly, the flanking regions of *ybbN* coding region was amplified by PCR using oligonucleotides listed in the Supplementary Table 1. After, these two fragments were cloned in pEX18Ap [18]. *P. aeruginosa* UCBPP-PA14 strains was then transformed with the resulting construct (named as pEX18Ap_del_*ybbN*) by conjugation with *E.coli* S17-1 strain harboring the pEX18Ap_del_*ybbN* construct. The *P. aeruginosa* transconjugants were first selected in LB plates supplemented with carbenicillin and after a second event of recombination, we selected only colonies that had lost carbenicillin resistance and grew in the presence of 10 % of sucrose (the product of *sacB* gene present in the pEX18Ap_del_*ybbN* construct that makes sucrose toxic to cells). Finally, these carbenicillin sensitive and sucrose resistant colonies were submitted PCR screening in order to identify the correct clones, i. e., those that presented a deletion in *ybbN* coding region.

### 2.2. Cloning, expression, and purification of recombinant YbbN enzymes from *X. fastidiosa* (*Xf*YbbN) and from *P. aeruginosa* (*Pa*YbbN)

The *XfybbN* (XF_2174) and *PaybbN* (PA14_11340) coding regions were PCR amplified using genomic DNA from the respective strains as template and the primers listed in Supplementary Table 1. The fragments were cloned into *Nde*I/*Xho*I restriction sites of pET28a expression vector (Novagen EMD Biosciences, Inc., Merck) or *Nde*I/*Bam*HI restriction sites of pET15b expression vector (Novagen EMD Biosciences, Inc., Merck), respectively. To express the recombinant proteins, overnight cultures of *E. coli* BL21(*DE3*) harboring pET28a-*XfybbN* or pET15b-*PaybbN* were used to inoculate 1 liter of fresh LB medium supplemented with appropriated antibiotics and cultured further until the OD_600nm_ reached 0.6– 0.8. The expression was induced by 0.5 mM isopropyl 1-thio-D-galactopyranoside (IPTG) at 20 °C. After overnight incubation, cells were harvested by centrifugation, suspended in the start buffer (50 mM Tris-HCl buffer, pH 8.0, containing 0.5 M sodium chloride) supplemented with lysozyme (100 μg/mL) and protease inhibitors (SIGMA # S8830) and disrupted by sonication under ice bath. After cell lysis, nucleic acids were precipitated with 1% streptomycin sulfate treatment and centrifuged during 40 min, 15000 rpm at 4 °C. The supernatant was loaded onto a nickel affinity column (Ni-NTA superflow columns from Qiagen) that was previously equilibrated with start buffer. To remove imidazole from His-YbbN containing fractions, the samples were submitted to a buffer exchange using HiTrap desalting columns (GE Healthcare). The recombinant proteins were submitted to an additional anion exchange chromatography step, using MonoQ 10/100 GL (GE HealthCare Inc.) column in order to obtain a pure fraction of *Xf*YbbN, *Ec*YbbN or *Pa*YbbN. Purified proteins were stored in 50 mM Tris-HCl buffer (pH 8.0) and 20 mM sodium chloride. Protein concentrations were determined spectrophotometrically at *A*_280 nm_.using the following extinction coefficients: (*Xf*YbbN, ∊_red_=22460 M^−1^.cm^−1^ and ∊_ox_=22710 M^−1^.cm^−1^) and *Pa*YbbN (∊_ox_ = 24660 M^−1^.cm^−1^ and ∊_red_ = 24410 M^−1^.cm^−1^) that were calculated using the ProtParam tool (https://web.expasy.org/protparam/). *Xf*TrxR, *Xf*TsnC, *Xf*Ohr, *Xf*PrxQ used in this work were expressed and purified as previously described [19,20].

### 2.3. Analysis of *Xf*YbbN and *Ec*YbbN interaction with *Ec*GroEL

For the pull-down assay, *E.coli* BW25113 cells were collected at exponential phase of growth in LB medium, resuspended in 50 mM Tris-HCl pH 8.0 buffer and disrupted on ice by sonication. As bait proteins, purified His-tagged *Ec*YbbN and *Xf*YbbN were immobilized in a Ni-NTA-agarose resin and washed three times with a 50 mM Tris-HCl pH 8.0 buffer. Then, bait proteins bound to Ni^2+−^ agarose resin were incubated with *E.coli* whole cell lysate for 1 hour at 4 °C under gently agitation. After that, each bait protein bound to the Ni-NTA-agarose resin was washed four times with cold 50 mM Tris-HCl pH 8.0 buffer. To elute protein complexes from Ni^2+−^ agarose resin, reducing SDS-PAGE sample buffer [21] was added directly to the resin and the protein complexes were resolved by SDS-PAGE.

As negative controls, unbounded Ni-NTA-agarose resin or *Pa*Ohr bound to Ni-NTA-agarose resin was incubated with *E.coli* whole cell lysate in the same way that was described above for bait loaded resin.

To perform Western Blot assays, protein complexes resolved by SDS-PAGE were transferred to nitrocellulose membrane and blocked with 5% non-fat milk in TBS buffer for 30 minutes. After, membrane was incubated with polyclonal rabbit antisera anti-GroEL from *Caulobacter crescentus* [22] at 1:5000 dilution in TBS 5% non-fat milk buffer during 1 hour at room temperature, washed and incubated with HRP-conjugate secondary anti-rabbit monoclonal antibody (Cell Signaling #7074) at 1:10000 dilution in TBS buffer for 1 hour. The immune complexes were revealed with the chemiluminescence reagent ECL Prime (GE Healthcare, UK).

### 2.4. DNA site-directed mutagenesis

Site-directed mutagenesis was performed according to the manufacturer’s instructions of QuikChange® Site-Directed Mutagenesis Kit (Agilent Technologies). The reactions were conducted using as template the plasmid pET28a-*XfybbN* and pET15b-*Ec*YbbN, carrying the wild-type *ybbN* from *X. fastidiosa* and *E.coli*, respectively. The pairs of oligonucleotides utilized to generate the *Xf*YbbN C37A and C40A and *Ec*YbbN S35C substitutions are listed in Supplementary Table 1. The mutations were confirmed by sequencing and the result constructs were named as pET28a-*Xfybbn_C37A* and pET28a-*Xfybbn_C40A* and pET15b-*Ecybb*N_*S35C*.

### 2.5 Disulfide reductase activity assays

#### 2.5.1 Insulin reduction assay

A fresh solution of insulin diluted in 100 mM of PBS buffer was added to a final concentration of 0.13 mM in a mix containing 0.1 M of sodium phosphate pH 7.0, 2 mM of EDTA, 0.33 mM of DTT and 10 μM of YbbN. The light scattering due to insulin aggregation was followed at 650 nm at room temperature. As positive controls, recombinant Trx from *X. fastidiosa* (*Xf*TsnC), *P. aeruginosa* (*Pa*TrxA) or *E.coli* (*Ec*Trx) were analyzed at 10 μM concentration. Addition of DTT alone, without any protein, served as negative controls. The assay was performed in a 96 well plate at the Synergy H1 Microplate Reader, BioTek Instruments, Inc.

#### 2.5.2 DTNB assay

DTNB (Ellman’s reagent) was used to measure disulfide reductase activities. The reaction was carried out in 20 mM Tris-HCl (pH 8.0) buffer, 100 μM DTPA, 500 μM DTNB (Sigma-Aldrich, St. Louis, USA), 200 μM NADPH (Sigma-Aldrich, St. Louis, USA), 100 nM *Xf*TrxR and started with the addition of 2 μM YbbN. The reduction of DTNB to TNB was followed at 412 nm in a 96 well plate using Synergy H1 Microplate Reader, BioTek Instruments, Inc. In order to compare the rates of DTNB reduction by *Xf*YbbN or by *Pa*YbbN, the same assay was also conducted using recombinant Trx from *X. fastidiosa* (*Xf*TsnC) or from *P. aeruginosa* (*Pa*TrxA).

#### 2.5.3 Trx System coupled Assay

To test if the peroxidase activity of *Xf*PrxQ can be supported by *Pa*YbbN or by *Xf*YbbN, we followed NADPH consumption (∊_340nm_ = 6220 M^−1^.cm^−1^) coupled to the corresponding Trx System. *Xf*YbbN or *Pa*YbbN (1 μM) were incubated with *Xf*PrxQ (1 μM), *Pa*TrR (PA14_53290) or *Xf*TrR (XF_1448) (0.2 μM) and NADPH (200 μM) in 1 mL of sodium phosphate (50 mM) buffer, pH 7.4 at 37 °C. After temperature equilibration, reactions were started by H_2_O_2_ (200 μM) addition.

### 2.6. Evaluation of HOCl-activated holdase activity

For all assays, the concentration of sodium hypochlorite solution (SIGMA #425044) was ascertain spectrophotometrically by measuring its absorbance in 200 mM potassium hydroxide at pH 12 (∊ ^292nm^ = 350 M^−1^cm^−1^) [23].

*Xf*YbbN, *Pa*YbbN or *Ec*YbbN were treated with ten-fold molar excess of HOCl for 15 minutes at room temperature. The excess of HOCl was removed by gel filtration and treated proteins used in the follow assays:

#### 2.6.1 Nile Red assay

Nile Red (SIGMA #72485) was used to assess the changes in the hydrophobicity of YbbN proteins. The experiments was performed as described in [24]. Each assay was performed in triplicate from two independent batches of purified proteins.

#### 2.6.2 HOCl-activated holdase activity assay

The ability of HOCl-treated *Xf*YbbN, *Pa*YbbN or *Ec*YbbN to inhibit the thermal aggregation (43 °C) of citrate synthase (SIGMA #C3260) was compared with the protein without treatment. All assays were carried out at 3YbbN:1CS molar ratio. Each assay was performed in triplicate from two independent batches of purified proteins.

### 2.7. Functional categorization of YbbN proteins

The *Xf*YbbN amino acid sequence was used as query to perform an iterative profile search using HmmerWeb version 2.38.0 [25] against UniProtKB database in 06/25/2019. All proteins that did not present the same domain architecture of *Xf*YbbN were discarded. From Jackhmmer output file, we created a set of representative sequences with an identity cut-off of 95% using CD-HIT software [26]. The sequence alignment was performed using MAFFT [27] and gaps present in more than 90 % of aligned sequences were removed to build a profile HMM and create a HMM logo with Skylign [28].

### 2.8. Thermal, HOCl and organic hydroperoxides sensibility assays

Growth curves of wild type and Δ*ybbN P. aeruginosa* strains in the presence of increasing concentrations of HOCl were analyzed. *P. aeruginosa* was grown on liquid LB medium for 16 hours, 240 rpm at 37 °C. Cells were washed in M9 + glucose and diluted in the same minimal medium to an OD_600nm_ = 0.05 supplemented with incremental amounts of HOCl. We followed the growth for 16 hours.

The sensitivity of strains to a several stressors were measured by a modified Kirby-Bauer test. Strains were grown on LB medium until reach OD_600nm_ = 0.1 and 100 μL of cell suspension were spread in solid medium and let to air dry. After that, 6 mm paper filter disks were saturated with 10 μL of HOCl (500 mM), paraquat (1 M), diamide (2 M) or DTT (4 M) solutions and placed on the surface of inoculated LB plates. After 16 hours, the size of inhibition zone was measured.

## 3. Results

### 3.1. Functional categorization of YbbN proteins considering the presence or absence of conserved Cys residues in the Trx domain

We were interested to investigate the sequence variations among YbbN proteins, considering the composition of the C/SXXC/S motif and the presence or absence of a Cys residue equivalent to Cys 63 in *Ec*YbbN of Trx domain [11,29]. Our search retrieved 11942 sequences that presented the YbbN architecture, i.e., N-terminal Trx-domain fused with C-terminal TRP-domain. We built a profile hidden Markov model (HMM) to represent the multiple sequence alignment of our analyses (Fig. 1A). Proteins with Trx and TPR domains in a distinct arrangement were not considered here. For instance, AtTDX is a protein from *Arabidopsis thaliana* that also have a multidomain structure formed by a Trx-domain linked to a TPR-domain, but the Trx-domain are located at C-terminal portion and the TPR-domain at the N-terminal [30], was not considered here.

**Figure 1.**
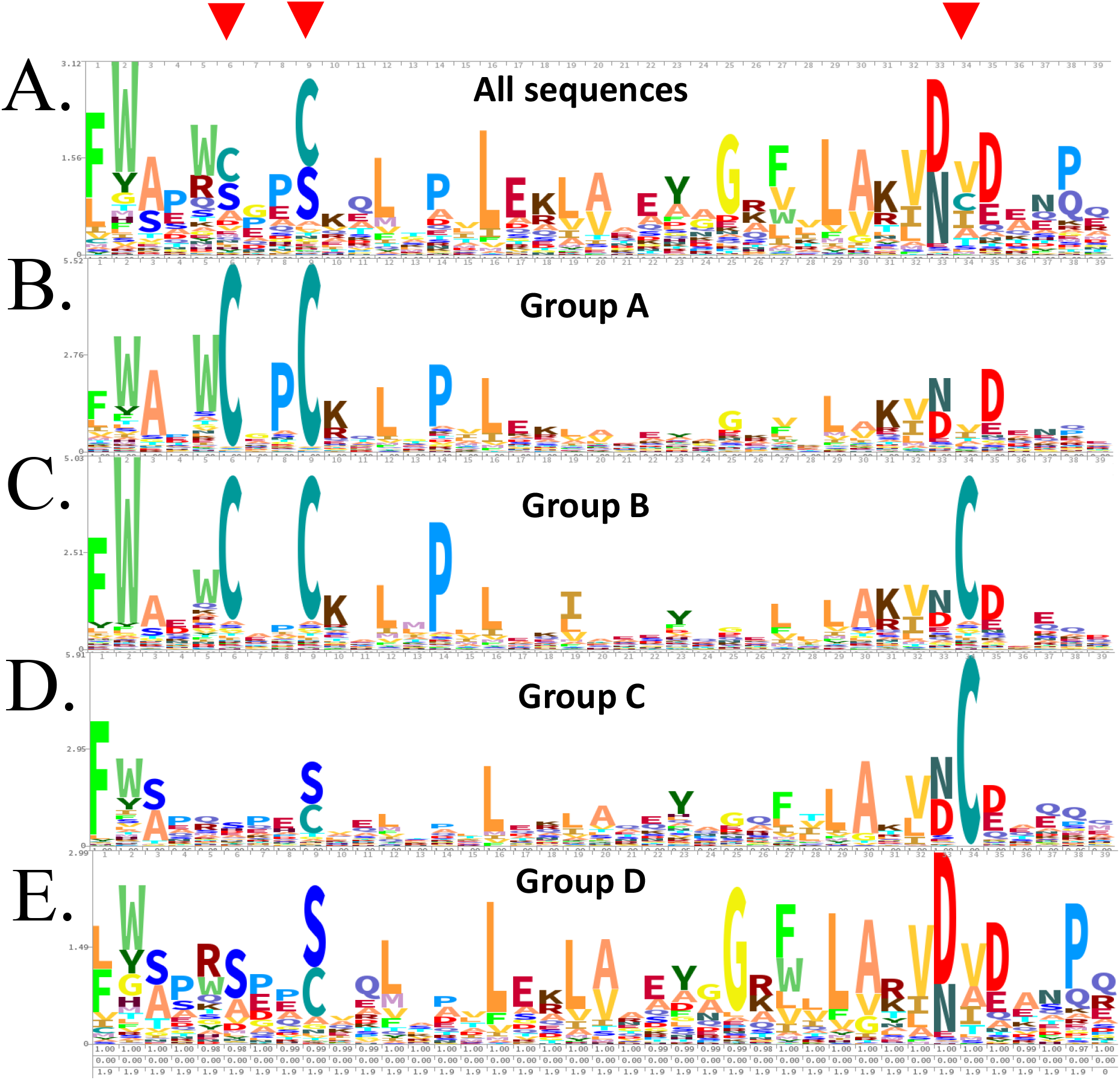
Graphical representation of CXXC[N24]C motif from *Xf*YbbN/CnoX homologue sequences. Multiple sequence alignment was used to produce a profile hidden Markov model (profile HMM). All profile HMM were visualized by HMM logo created with Skylign. **(A)** HMM logo of all 5532 representative sequences (95 % identity cut-off) extracted from UniProtKB database. **(B)** HMM logo of 3768 sequences representative of *Xf*YbbN group A (CXXC[N_24_]X). In this group, CXXC motif is fully conserved but lacks the additional Cys residue. **(C)** HMM logo of 219 sequences representative of *Pa*YbbN group B (CXXC[N_24_]C). In this group, CXXC motif is fully conserved and the additional Cys residue are fully conserved. **(D)** HMM logo of 537 sequences representative of *Ec*YbbN group C (SXXC[N_24_]C). In this group, the CXXC motif is degenerated and the additional Cys is fully conserved. (**E)** HMM logo of 1009 sequences representative of a group D that have CXXC motif is degenerated and lacks the additional Cys residue.

Based on this analysis, we propose that the YbbN family of proteins should be functionally divided in four groups: group A (CXXC[N_24_]X), group B (CXXC[N_24_]C), group C (XXXC/S[N_24_]C) and group D (XXXC/S[N_24_]X). We then created an HMM profile for each one of these groups (Fig. 1 B-E).

Accordingly, most of the proteins belongs to the groups A and B (68 % - 3987 of 5533), presenting the CXXC motif, as observed in *Xf*YbbN and *Pa*YbbN (Fig. 1 B and 1 C, respectively). Members of groups A and B were separated due to the presence or absence of Cys residues equivalent to Cys63 of *Ec*YbbN.

In contrast, members of groups C and D do not present the two Cys in positions equivalent to the CXXC motif (Fig. 1 D and 1 E). Noteworthy, most of these sequences do not display a single conserved Cys residue in this motif (Fig. 1 D and 1 E). As in the case of groups A and B, members of group C and D were also separated due to the presence or absence of a Cys residue equivalent to Cys 63 of *Ec*YbbN (Fig. 1 D and 1 E). Therefore, according to this proposal, *Ec*YbbN that contains a SQHC motif and Cys63 is a member of the group C.

This analysis allowed us to identify two new groups of YbbN proteins: one comprising enzymes that conserve both CXXC motif and a Cys63 equivalent (group B) and other one that do not display neither a conserved CXXC motif nor a *Ec*YbbN Cys63 equivalent (group D).

In group D, it is interesting to note that the lack of a conserved CXXC motif is accompanied by a lack of other conserved residues present in Trx family (compared to Pfam [25] Trx HMM logo - accession code PF00085) that could reflect the absence of disulfide reductase activity, as observed in *Ec*YbbN [29].

In contrast, group B present HMM profile quite similar to the group A (compare Fig. 1 B and C), therefore, it is reasonable to assume that these proteins are also endowed with disulfide reductase activity. It is also interesting that members of this group are restricted mainly to Pseudomonadales and Alteromonadales from Gammaproteobacteria class, which includes some relevant opportunistic pathogens like *P. aeruginosa*. Next, we decided to compare proteins belonging to the groups A, B and C, represented by *Xf*YbbN, *Pa*YbbN and *Ec*YbbN, respectively.

### 3.2. *Xf*YbbN and *Pa*YbbN are functional disulfide reductases

Since both *Xf*YbbN and *Pa*YbbN contain the two Cys residues in the CXXC motif as the canonical Trx enzymes, we investigated whether these two proteins are endowed with disulfide reductase activity.

Initially, we observed that *Xf*YbbN and *Pa*YbbN accelerated the reduction of insulin disulfide bonds by DTT, which is a classical assay of Trx enzymes (Fig. 2 A and C, respectively). Noteworthy, the insulin reductase activities of YbbN proteins were comparable to the classical Trx (*Xf*TsnC and *Pa*Trx) activities, which were also analyzed as positive controls (Fig. 2 A and C). As expected, *Ec*YbbN did not reduce insulin, although *Ec*Trx did (Fig. 2 B).

**Figure 2.**
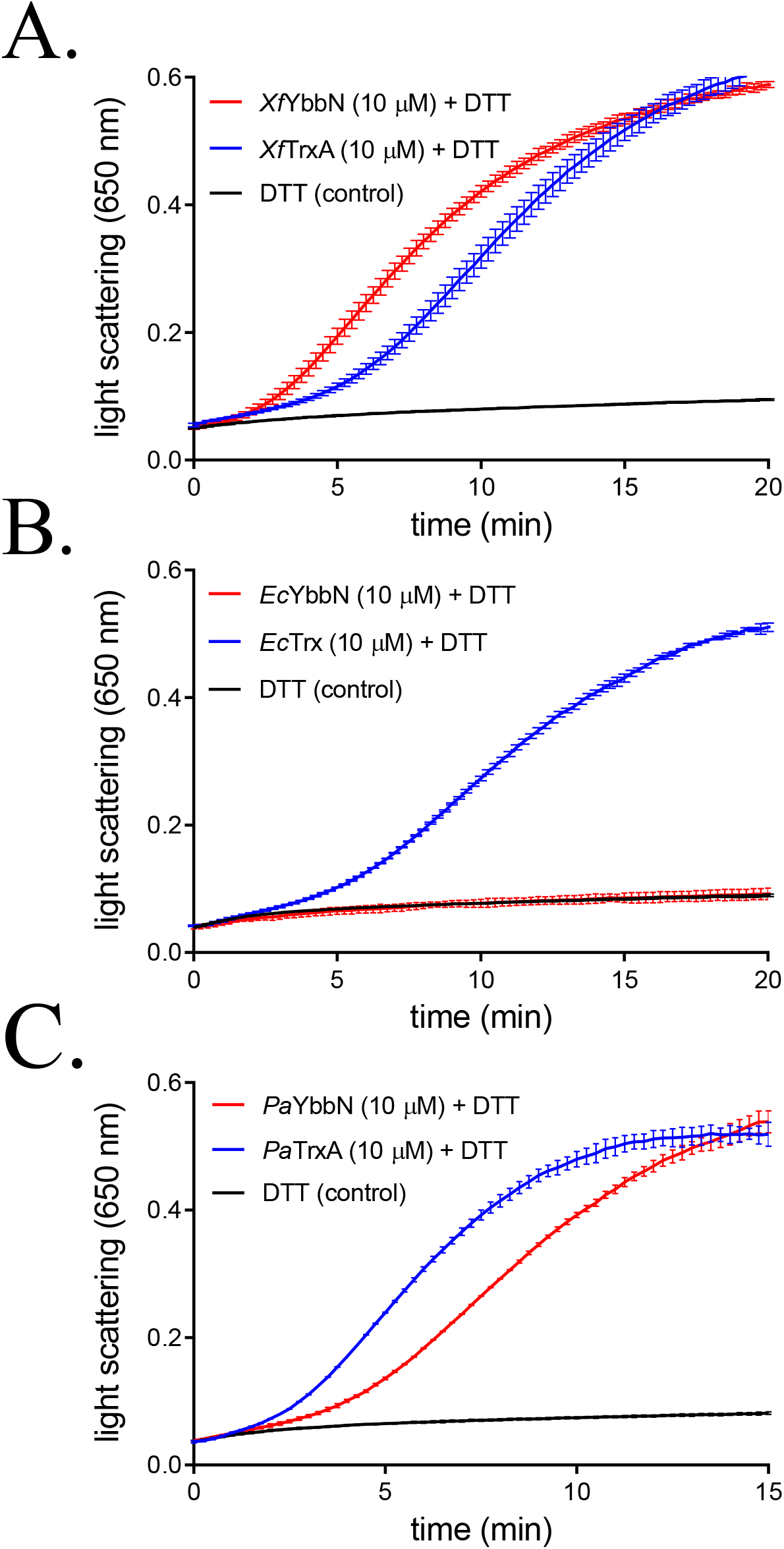
Insulin reductase activities of recombinant YbbN from *X. fastidiosa* and *P. aeruginosa*. The protocol was adapted to be performed in a 96 well plate. In the assay, a fresh made solution of insulin diluted in 100 mM of PBS buffer was added to a final concentration of 0.13 mM in a mix containing 0.1 M of sodium phosphate pH 7.0, 2 mM of EDTA, 0.33 mM of DTT and 10 μM of *Xf*YbbN **(A)**, *Ec*YbbN **(B)** or *Pa*YbbN **(C)**. The light scattering due aggregation of insulin was followed at 650 nm at room temperature. Each assay includes native Trxs (10 μM of recombinant *Xf*Trx (TsnC), *Pa*TrxA or *Ec*Trx) as positive controls. Reaction tubes containing DTT and insulin was the negative control (black line of each graph). Each assay was performed in triplicate and the presented data is representative of results obtained from two independent batches of purified proteins.

Since insulin reduction is a qualitative assay, we also analyzed the reduction of DTNB (an artificial disulfide) by these oxidoreductases in the presence of their cognate thioredoxin reductase (TrxR) and NADPH.

*Xf*YbbN reduced DTNB with high efficiency, again comparable with the reduction of DTNB by *Xf*TsnC (Fig. 3 A and B). Indeed, the K_m_ values for the reduction of *Xf*TsnC or *Xf*YbbN by *Xf*TrxR were very similar (Table 1).

**Figure 3.**
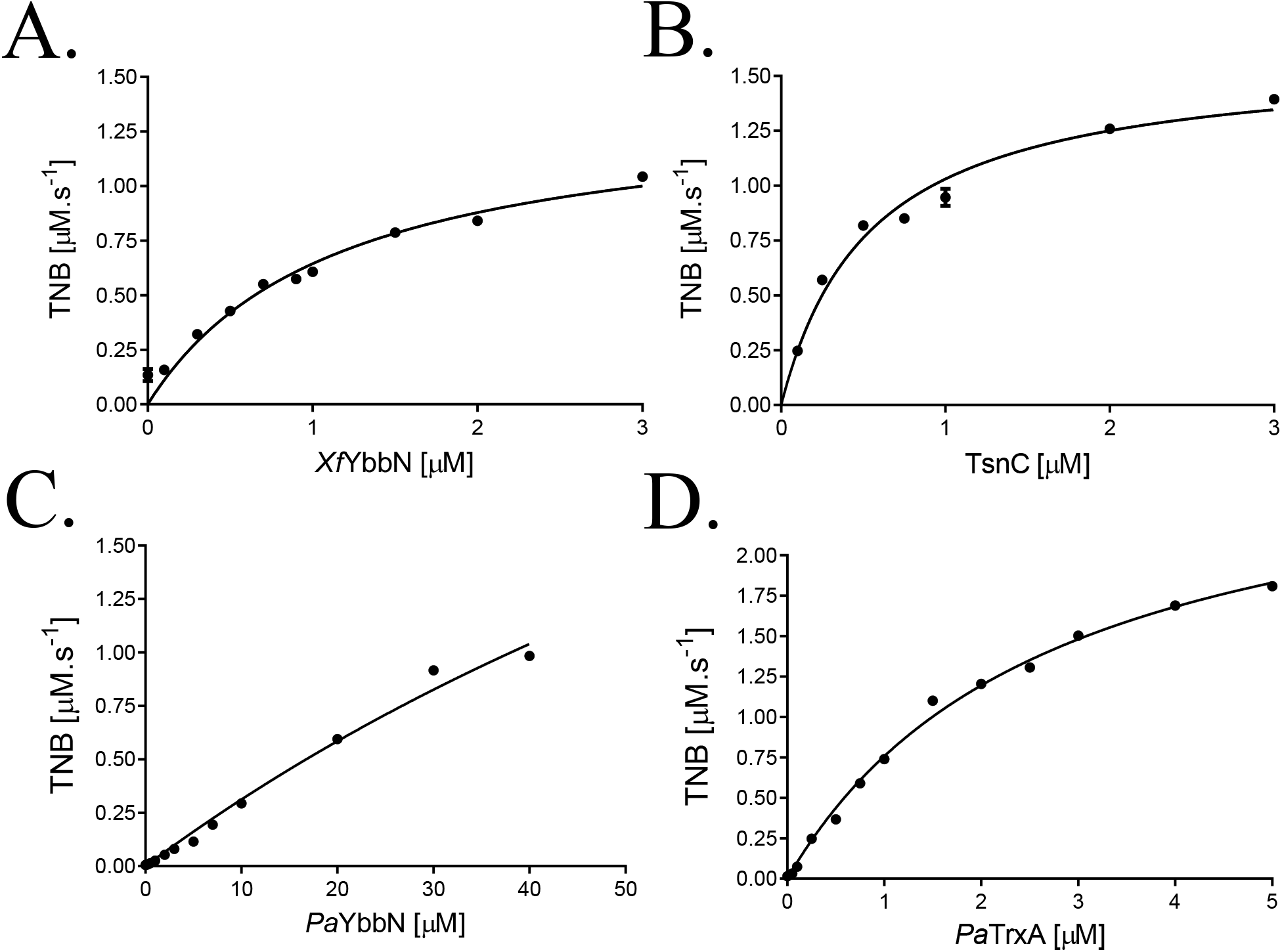
*Xf*YbbN and *Pa*YbbN are functional disulfide reductase. Assays were carried out in Tris-HCl 20 mM pH 8.0, and included DTPA 100 μM, DTNB 500 μM, *Xf*TrxR (A and B) or *Pa*TrxR 0.1 μM (C and D) and NADPH 200 μM. Rates of TNB formation in the presence of variable enzyme concentrations, *Xf*YbbN (A), TsnC (B), *Pa*YbbN (C) or *Pa*Trx (D), were analyzed by non-linear regression fitted to the Michaelis-Menten equation.

**Table 1.**
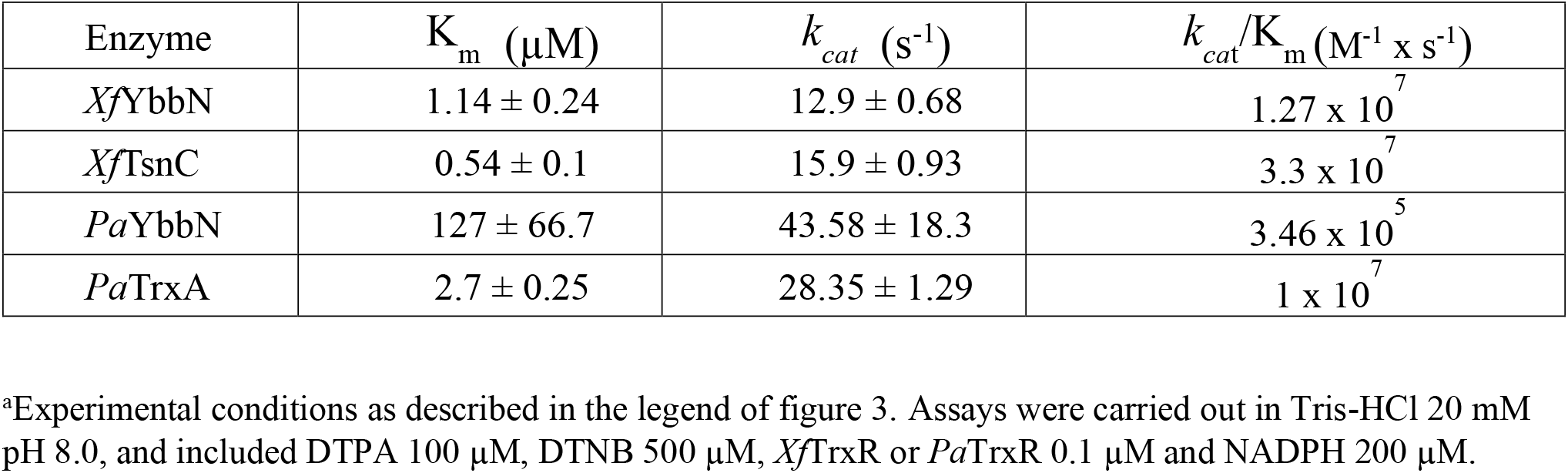
Apparent kinetic parameters of YbbN and Trx enzymes as substrates of their cognate TrxR enzymes^a^.

In contrast, *Pa*YbbN was reduced by NADPH/ PaTrxR, but the catalytic efficiency was lower than the similar reduction of *Pa*Trx (Fig. 3 C and D). Indeed, the K_m_ values of *Pa*YbbN is two order of magnitude higher than the K_m_ values of *Pa*Trx (Table 1), indicating that the latter is a better substrate for *Pa*TrxR. YbbN from a nonpathogenic freshwater living bacteria *Caulobacter crescentus* (*Cc*YbbN) also displayed DTNB reductase activity and it is reduced by TrxR from *C. crescentus* with similar efficiency as cognate Trx [29]. According to our analysis, *Cc*YbbN belongs to the same group of *Xf*YbbN (group A).

### 3.3. *Xf*YbbN and *Pa*YbbN are substrates for Cys-based peroxidases

We also investigated if *Xf*YbbN might also reduce disulfides in Cys-based peroxidases from *X. fastidiosa*, as classical Trx enzymes are well known to support the peroxidase activities of peroxiredoxins. Two thiol peroxidases were investigated: *Xf*PrxQ, whose peroxidase activity is supported by *Xf*TsnC [19] and the organic hydroperoxide resistance protein (Ohr), that is not reduced by *Xf*TsnC [20].

*Xf*YbbN supported the peroxidase activity of *Xf*PrxQ (Fig 4 A and C) as well as *Xf*TsnC (Fig. 4 D), presenting K_m_ values of 2.55 ±1.01 (Fig. 4 C) and 1.96 ±0.74 μM (Fig. 4 D), respectively. In contrast, *Xf*YbbN did not support the *Xf*Ohr peroxidase activity (Supplementary Fig. 1).

**Figure 4.**
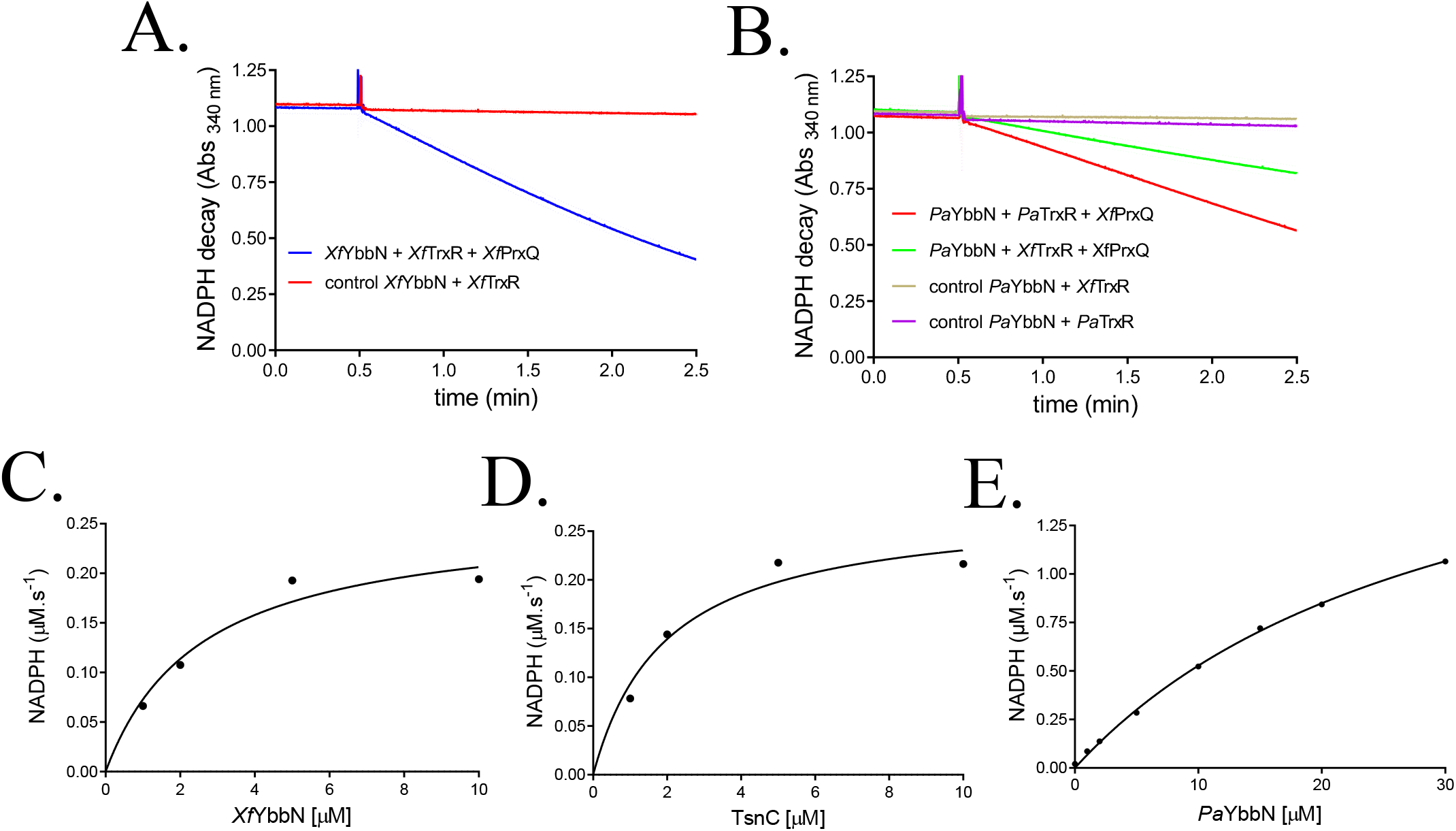
*Xf*PrxQ peroxidase activity is sustained by *Xf*YbbN and *Pa*YbbN. The coupled assays were carried out at 37 °C in sodium phosphate 50 mM pH 7.4 buffer, 100 μM DTPA, 1 μM *Xf*PrxQ, 1 μM *Xf*YbbN **(A)** or *Pa*YbbN **(B)**, 200 μM NADPH, 0.1 μM *Xf*TrxR or *Pa*TrxR and 0.2 mM t-BOOH, to start the reaction. Rates of NADPH oxidation were determined in 1-10 μM range for *Xf*YbbN **(C)**,*Xf*TsnC **(D)** or 0.25-40 μM range for *Pa*YbbN **(E)** at 37 °C in sodium phosphate 50 mM pH 7.4 buffer, 1 μM *Xf*PrxQ,. 0.2 μM *Xf*TrxR or *Pa*TrxR, 200 μM NADPH and 0.2 mM H^2^O^2^, to start the reaction. V_max_ and K_m_ values were obtained by non-linear regression fitted to the Michaelis-Menten equation.

In order to test if *Pa*YbbN could also support the activity of thiol peroxidases, we evaluated the ability of *Pa*YbbN to reduce *Xf*PrxQ (Fig. 4 B). *Pa*YbbN also sustained the activity of *Xf*PrxQ, however displaying a K_m_ value (31.26 ± 2.18 μM) ten-fold times higher (Fig. 4E).

Altogether, the above results suggested that YbbN enzymes containing the CXXC motif reduce disulfide bonds of biological substrates. Recently, it was also described that *Cc*YbbN supports the activities of methionine sulfoxide reductase and DsbDα [29].

### 3.4. *Xf*YbbN but not *Pa*YbbN displayed a potent holdase activity upon HOCl treatment

Recently, *Ec*YbbN was described as an efficient holdase upon HOCl treatment [11]. The chlorination of specific amino acids residues at the TPR domain turns the protein more hydrophobic, activating its holdase activity [11]. In contrast, CcYbbN displays a holdase activity independently of the HOCl treatment [29]. We also evaluated if *Xf*YbbN and *Pa*YbbN could also prevent Citrate Synthase (CS) from thermal aggregation. At 3:1 molar ratio, *Xf*YbbN was not able to prevent CS from thermal aggregation (Fig. 5 A, blue line). Notably, *Xf*YbbN displayed a holdase activity when pre-treated with HOCl (Fig. 5 A, red line). Therefore, *Xf*YbbN behave similarly to *Ec*YbbN [11], but differently than *Cc*YbbN [29]. As reported before [11], we also observed that *Ec*YbbN only presented a holdase activity upon HOCl treatment (Fig. 5 B). In contrast to *Ec*YbbN and *Xf*YbbN, *Pa*YbbN did not show an holdase activity, even after HOCl treatment (Fig. 5 C).

**Figure 5.**
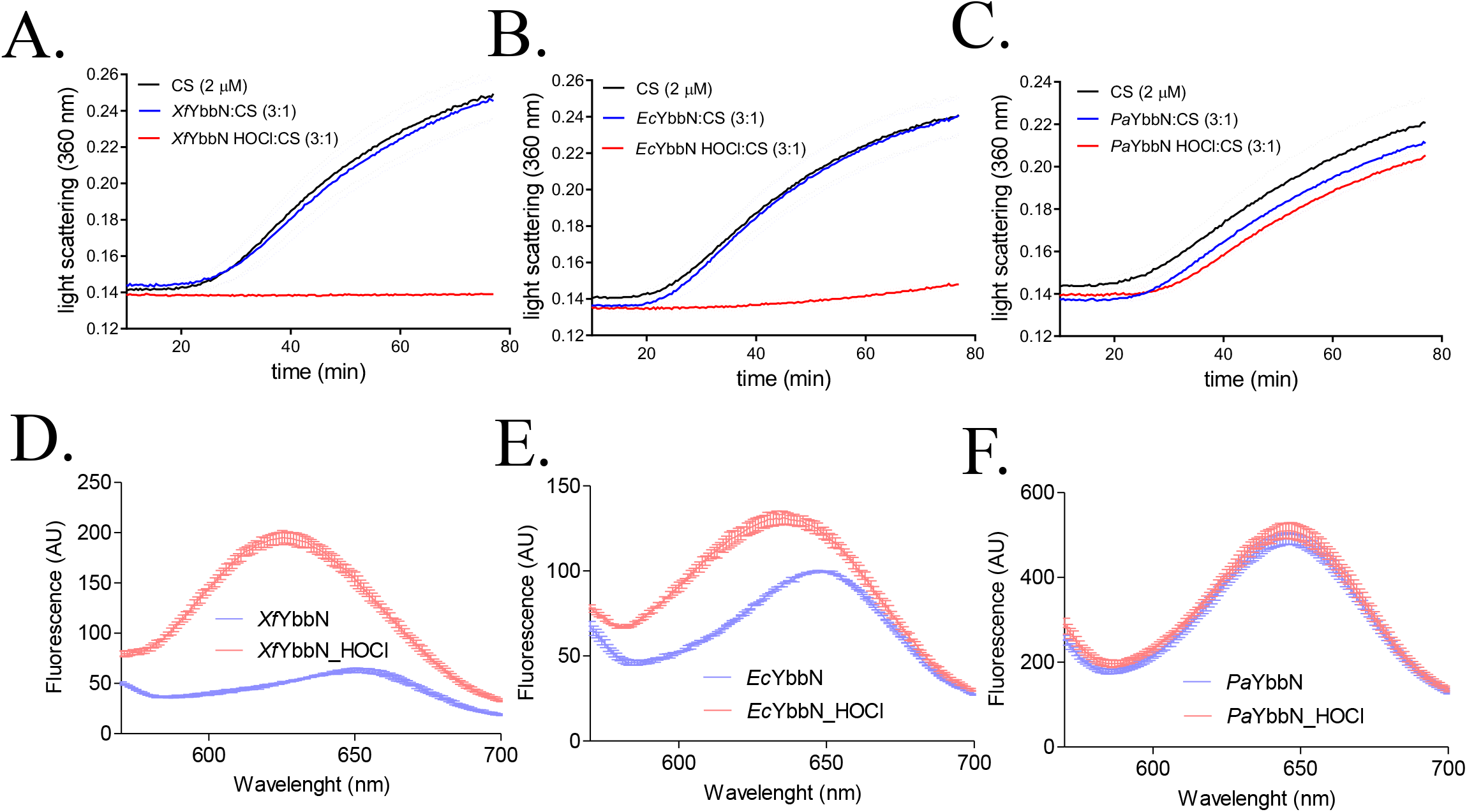
Chaperone holdase activity of HOCl-treated *Xf*YbbN, *Pa*YbbN and *Ec*YbbN. Each protein was treated with ten-fold molar excess of HOCl for 15 minutes at room temperature. After removal of HOCl by gel filtration, the ability of HOCl-treated *Xf*YbbN **(A)**, *Ec*YbbN (positive control) **(B)** or *Pa*YbbN **(C)** to inhibit the thermal aggregation (43 °C) of citrate synthase was compared with the protein without treatment. We also monitored the change in the Nile Red fluorescence in the presence of HOCl-treated *Xf*YbbN **(D)***Ec*YbbN (positive control) **(E)** or *Pa*YbbN **(F)** compared with Nile Red fluorescence in the presence of proteins without treatment. Each assay was performed in triplicate. Presented data is representative of results obtained from two independent batches of purified proteins.

We also monitored the polarity of YbbNs as a function of HOCl treatment by the Nile Red assay. Noteworthy, only the YbbN proteins with HOCl-induced holdase activity, i. e. *Xf*YbbN and *Ec*YbbN, presented an increase on its hydrophobicity (Fig. 5 D and E, respectively). The Nile Red fluorescence profile of *Pa*YbbN was nearly identical for the protein with or without HOCl treatment (Fig. 5 F). Therefore, the induction of the *Ec*YbbN and *Xf*YbbN holdase activities by HOCl involves increase hydrophobicity. In contrast, *Pa*YbbN does not display holdase activity and its hydrophobicity is not altered by HOCl treatment (Fig. 5). Therefore, the abilities of YbbN enzymes to display HOCl – activated holdase activities varied considerably and did not correlate with the presence of a CXXC or SXXC motif.

### 3.5. Effects of site-directed mutagenesis in CXXC/SXXC motifs of YbbN enzymes

Next, we investigated the effects of modifying the composition of SXXC or CXXC motifs on the oxidoreductase and holdase activities of YbbN enzymes. Initially, we analyzed if *Ec*YbbN would gain disulfide reductase activity by mutating the SQHC motif to CQHC (*Ec*YbbNS35C). Remarkably, the substitution of a single amino acid (*Ec*YbbNS35C) made this engineered enzyme capable to reduce the disulfides of insulin at rates comparable to the canonical Trx (*Ec*TrxA) (Fig. 6 A, red line). However, *Ec*YbbNS35C did not reduce DTNB using *Ec*TrxR/NADPH as the reducing power. Probably, *Ec*TrxR was not capable to interact properly with *Ec*YbbNS35C. Indeed, protein-protein interactions within components of the thioredoxin system are highly specific [31].

**Figure 6.**
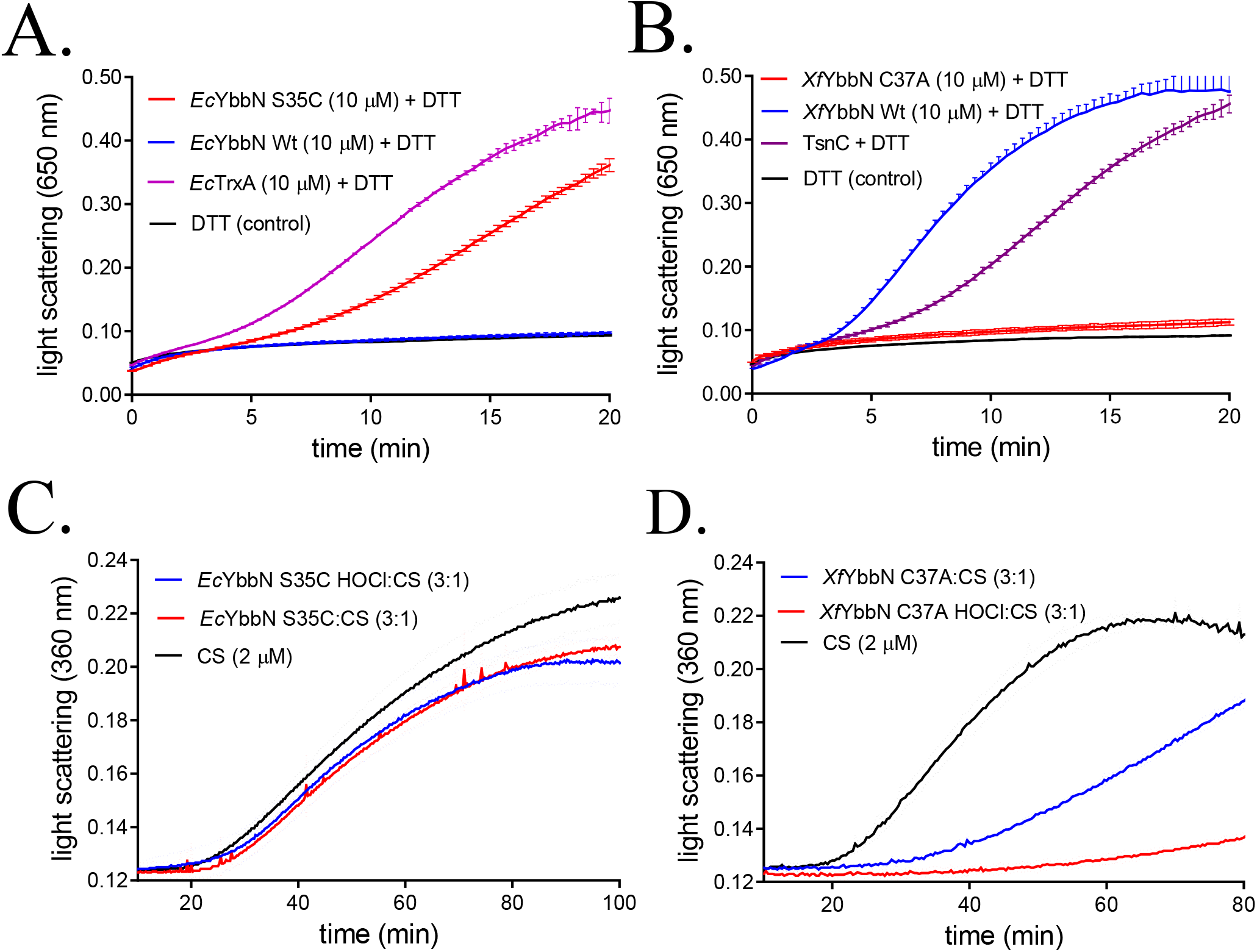
Oxidoreductase (A and B) and chaperone holdase activity (C and D) of HOCl-treated *Ec*YbbN S35C and *Xf*YbbN C37A. In the oxidoreductase assay, the light scattering due aggregation of insulin was followed at 650 nm at room temperature for *Ec*YbbN S35C **(A)** and *Xf*YbbN C37A **(B)**. For the graphs **(A)** and **(B)**, purple and blue lines represent insulin aggregation in the presence of 10 μM of recombinant thioredoxins or 10 μM of wild type YbbN (positive controls), respectively and the black line represent the reaction in a tube containing buffer plus DTT and insulin (negative control). In the chaperone assays, each YbbN protein was treated with ten-fold molar excess of HOCl for 15 minutes at room temperature. After removal of HOCl by gel filtration, the ability of HOCl-treated *Ec*YbbN S35C **(C)** or *Xf*YbbN C37A **(D)** to inhibit the thermal aggregation (43 °C) of citrate synthase (red line) was compared with the protein without treatment (blue line). For the graphs **(C)** and **(D)**, the black line represents the profile of CS thermal aggregation in the presence of buffer Each assay was performed in triplicate. Presented data is representative of results obtained from two independent batches of purified proteins.

Surprisingly, insertion of a Cys residue to the N-terminal position of SXXC motif (*Ec*YbbNS35C) abolished HOCl-induced holdase activity of *Ec*YbbN (Fig. 6 C). Noteworthy, in the same experimental conditions, wild type *Ec*YbbN displayed HOCl-induced holdase activity (Fig. 5 C). Therefore, while *Ec*YbbNS35C gained oxidoreductase activity (Fig. 6 A), it lost the HOCl-induced holdase activity (Fig. 6 C). These results suggest the oxidoreductase and the holdase activities of the YbbN enzymes are somehow interconnected. Noteworthy, it was reported before that mutation of the other Cys residue (*Ec*YbbNC38S) did not affect the HOCl-induced holdase activity [11].

We next evaluated the effects of replacing the N-terminal Cys residue by Ala of the CXXC motif of *Xf*YbbN (*Xf*YbbN C37A), thereby making this oxidoreductase more similar to *Ec*YbbN. Mutation of the Cys37 residue resulted in an enzyme without oxidoreductase activity (Fig. 6 B), while retaining its holdase activity upon HOCl treatment (Fig. 6 D). Noteworthy, the holdase activity of *Xf*YbbN C37A without treatment seemed to be higher than wild type *Xf*YbbN. Again, these results indicated some relationship between the holdase and oxidoreductase activities by still unknown mechanisms.

### 3.6 *P. aeruginosa* Δ*ybbN* strain did not exhibit increased sensitivity to heat shock, oxidative and HOCl stresses

No investigation of the roles of YbbN proteins containing the three conserved Cys residues (Group B, Fig. 1C) were performed in a cellular context. Therefore, we investigated *Pa*YbbN, as *P. aeruginosa* is exposed to HOCl during host-pathogen interactions [32,33].

The disruption of *ybbN* gene from *P. aeruginosa* did not render this bacteria strain more sensitive to diamide, paraquat, DTT (Supplementary Fig. 2 A-C, respectively) and to organic hydroperoxides (Supplementary Fig. 2 D and E). We also observed that wild type and Δ*ybbN* strains of *P. aeruginosa* have similar abilities to deal with HOCl stress (Fig. 7 A-D), consistent with the finding that *Pa*YbbN displayed no HOCl-induced holdase activity (Fig. 5 B). Furthermore, WT and Δ*ybbN* strains were equally resistant to heat shock (Fig. 7 E), differently from that observed for *E.coli* [10].

**Figure 7.**
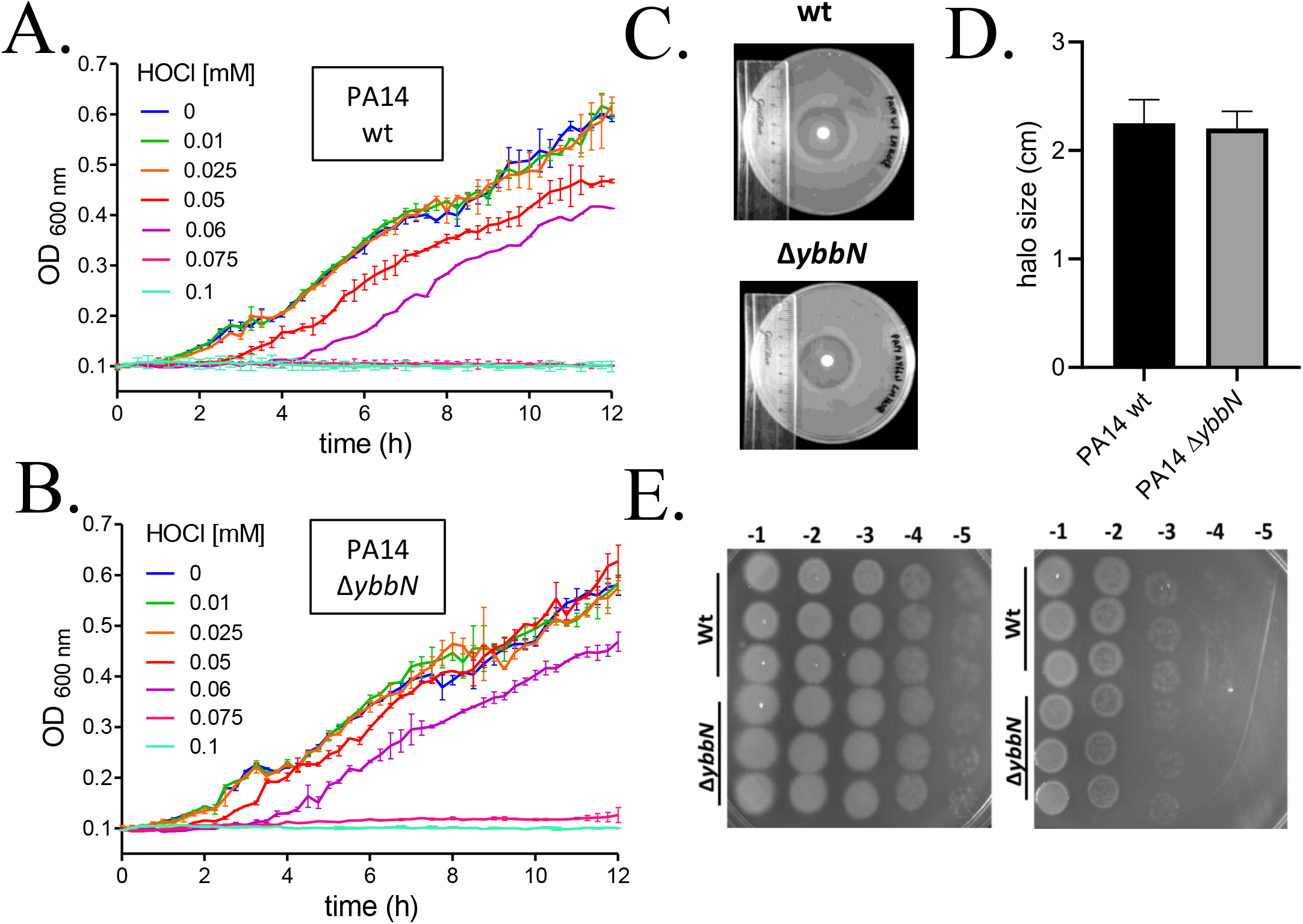
Effect of HOCl and heat shock treatment of PA14 wild type and Δ*ybbN* strains. **(A)** and **(B)** Growth curves of PA14 wild type and Δ*ybbN* strains, respectively, in the presence of increasing concentrations of HOCl. To perform the assay, saturated cultures of wild type and Δ*ybbN* strains were diluted in M9 medium supplemented with 0.2 % glucose to OD_600_ 0.1. After that, 180 μL of diluted cultures were distributed in separated wells of 96 well plates and incubated with increasing concentrations of HOCl. The final concentration of HOCl used is indicated in the graphs. Plates were incubated at 37 °C under constant agitation (282 rpm), and the growth profile was monitored during 12 h. Each growth curve on the graph represent the OD_600_ mean from three technical replicates. The experiment was repeated three times (three independent biological replicates)**. (C)** HOCl disk inhibition assay for wild type and *ybbN* mutant strains. **(D)** Statistical analysis of HOCl disk inhibition assay (*n*=24) showing that sensitivity to HOCl wt and *ybbN* mutant strains were not significantly different (P < 0.01). **(E)** Lack of YbbN does not affect sensitivity of *P. aeruginosa* cells to heat shock stress. The cultures were grown in LB medium at 37 °C to OD_600nm_ 0.5 and then shifted to 53 °C during 20 min. type and immediately before (time 0) and after 20 min, 10 μL of each dilution (10^−1^, 10^−2^ 10^−3^, 10^−4^ and 10^−5^) were spotted in LB plate and incubated at 37 °C for 16 hours.

Altogether, these results suggest that *Pa*YbbN does not play an essential role in the bacterial defense against HOCl, contrasting with the crucial role of *Ec*YbbN in similar process [11].

## 4. Discussion

YbbN proteins can present three activities: (i) holdase/co-chaperone; (ii) thiol-disulfide oxidoreductase and (iii) prevention hyperoxidation of target protein by formation of mixed disulfides [9,11,29]. The results described here indicated that the ability of each YbbN enzyme to display such activities depend on the C/S-X-X-C/S motif composition as well as on the presence or absence of a Cys residue homologous to Cys63 of *Ec*YbbN.

The composition of Cys residues in YbbN proteins ranges from proteins that do not contain any conserved Cys residues (Group D) to those that contain the two Cys residues in the CXXC motif plus a Cys residue homologous to Cys 63 of *Ec*YbbN (Group B). *Pa*YbbN is an example of an YbbN protein containing all the three conserved Cys residues and this work represents the first characterization of such an enzyme. Although at this point, it is still premature to make physiological considerations, it is important to observe that the biochemical properties of YbbN proteins vary a lot with the composition of the conserved Cys residues.

For instance, *Ec*YbbN does not present oxidoreductase activity and contain a SXXC motif. In contrast, *Xf*YbbN and *Pa*YbbN displayed disulfide reductase activity (Fig. 2, Fig. 3). As expected, the presence of a CXXC motif is an essential requirement for such activity. *Cc*YbbN also contains the CXXC motif and is endowed with thiol-disulfide oxidoreductase activity [29]. Notably, engineering *Ec*YbbN to present the CXXC motif, endowed this protein with disulfide reductase activity (Fig. 6 A).

The involvement of YbbN enzymes with the holdase/chaperone machinery was the first role described to these enzymes. Indeed, *Ec*YbbN co-purifies with GroEL, indicating strong physical interaction between these two proteins [9]. We also observed that *Xf*YbbN co-purified with GroEL from *E. coli* in nickel affinity chromatography (Supplementary Fig. 3 A) and by pull-down assays (Supplementary Fig. 3 B and C). *Ec*YbbN is a co-chaperone that is a negative regulator of GroEL in *E. coli* [9], although upon HOCl treatment this protein cooperates with GroEL/ES and DnaK/J/GrpE in the refolding of target proteins [11]. Furthermore, *Ec*YbbN is itself endowed with a holdase activity that is activated by the chlorination of some specific residues in the TPR domain [11]. Notably, insertion of a Cys residue in the SXXC motif of *Ec*YbbN abolished its holdase activity (Fig. 6 B), indicating that some redox reaction underlies this mechanism.

We also tried to identify a conservative pattern taking into account previously identified chlorinated residues in *Ec*YbbN: Y183, R182, H227, H242 and R276 [11]. However, multiple sequence alignment revealed that only R276 among these six residues is conserved among all YbbN protein sequences tested (Supplementary Fig. 4).

Furthermore, *Ec*YbbN is endowed with the ability to protect client proteins from irreversible oxidation and aggregation, by the formation of mixed disulfide bonds [11]. *Ec*YbbN prevents the hyperoxidation of other proteins by forming mixed disulfides through its Cys63 residue [11]. We demonstrated here that *Xf*YbbN and *Pa*YbbN present an alternative redox mechanism: support the peroxidase activity of *Xf*PrxQ. Therefore, *Xf*YbbN and *Pa*YbbN present similar roles as *Cc*YbbN, all them provide equivalents to reduce disulfide in Cys-based proteins [29].

HOCl production by phagocytic cells is one of the central mechanisms to protect host organisms from pathogenic bacteria, including *E. coli* and *P. aeruginosa* [32–34]. In turn, bacteria have to deal with this intense oxidative insult, with the involvement of Cys-based proteins [35–37]. Recently, it was described that *Ec*YbbN plays a central role in the *E. coli* response to HOCl [11]. However, wild type and Δ*ybbN* strains of *P. aeruginosa* were equally sensitive to HOCl stress (Fig. 7) and *Pa*YbbN had no holdase activity even when treated with HOCl (Fig. 5 C). These results suggest that *Pa*YbbN has a different role than *Ec*YbbN.

However, in our hands, both MG1655 and BW25113 *E.coli* Δ*ybbN* strains were equally resistant to HOCl as respective wild type strain (Supplementary Fig. 5 and 6, respectively), contrary to previously reported findings [11]. Therefore, more investigations are required to understand the roles of YbbN enzymes in bacterial response to HOCl. At this point, we do not understand the reasons for these contradictory results.

## Abbreviations

TPR: tetratricopeptide repeat
NADPH: nicotinamide adenine dinucleotide phosphate
Trx: thioredoxin
DTNB: 5,5-dithio-bis-(2-nitrobenzoic acid)

## Acknowledgments

We thank prof. Regina L. Baldini and Suely L. Gomes from Instituto de Química/Universidade de Sao Paulo/Brazil for the anti-GroEL antiserum and prof. Jean-François Collet from Université catholique de Louvain/Brussels for the *E.coli* K12 MG1655 wild type and derivative strains used in this study.

