## Supplementary data for "Investigation on the requirements for YbbN/CnoX displaying thiol-disulfide oxidoreductase and chaperone activities"

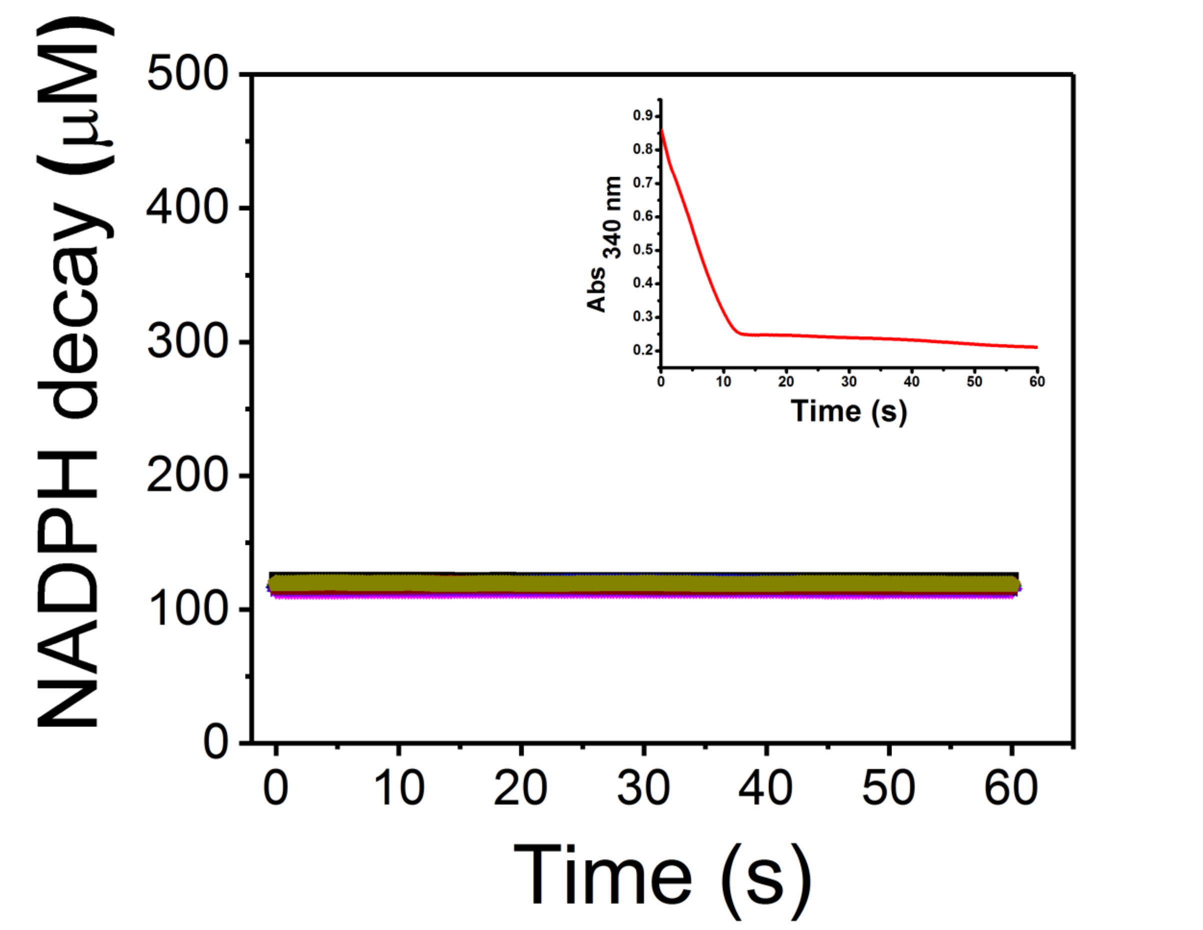


Supplementary Fig. 1. **Peroxidatic activity of *X. fastidiosa* Ohr linked to oxidation of NADPH by thioredoxin reductase/*Xf*YbbN system**. t-BOOH was used to start the reactions in a range of 10-200 μM. Reactions were carried out in Tris-HCl 20 mM pH 8.0 buffer, in the presence of NAPDH 150 μM, TrR 100 nM, *Xf*YbbN 3 μM and Ohr 1 μM. The inset represents a positive control of *Xf*Ohr with the LPD/lipoamide system to show that the enzyme is active and the reaction mixture in sodium phosphate buffer (50 mM), pH = 7.4 contained DTPA 0.1 mM, NADH 0.2 mM, lipoamide dehydrogenase from *X. fastidiosa* 1 uM, lipoamida 10 μM, Ohr 0.5 μM and t-BOOH 100 μM.

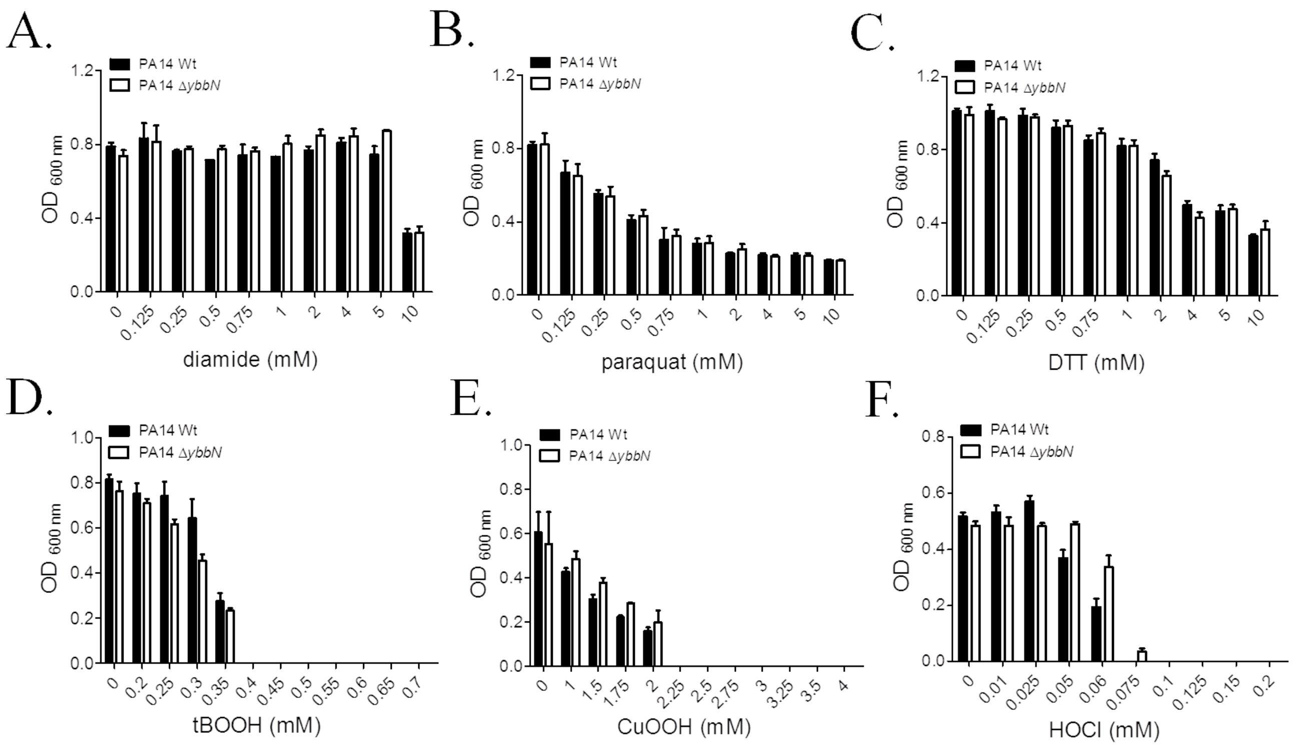


Supplementary Fig. 2. **Effect diamide (A), paraquat (B), DTT (C), t-BOOH (D), CuOOH (E) HOCl (F) treatment on the growth of PA14 wild type and Δ*ybbN* cultures.** To perform the assay, 180 µL of diluted cultures (final $OD_{600nm}$ 0.05) in LB (**A**), (**B**), (**C**), (**D**) and (**E**) or M9 (**F**) medium were distributed in separated wells of 96 well plates and incubated with increasing concentrations each stressor. The final concentration of each compound is indicated in the graphs. Plates were incubated at 37 °C under constant agitation and end-point $OD_{600nm}$ measured after 15 h of incubation. Each graph represent the media of three technical replicates from two independent biological replicates.

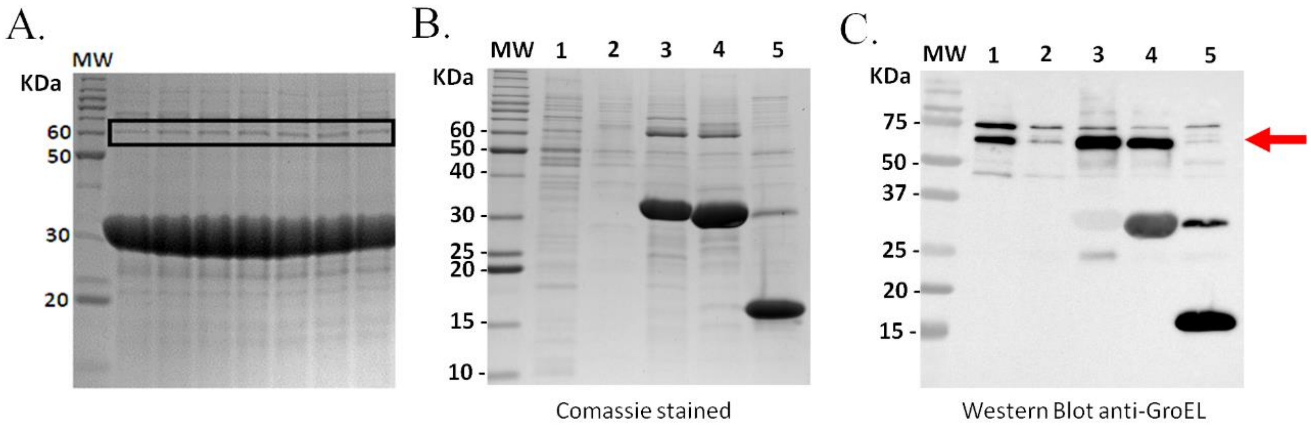


Supplementary Fig. 3. **Interaction of YbbNs with EcGroEL A.** *Xf*YbbN and *Ec*GroEL co-purify during Ni-NTA-agarose affinity purification. **B. and C.** Ni-NTA-agarose pull down assay showing that *Ec*YbbN and *Xf*YbbN interact with *Ec*GroEL. Lane 1 – *E.coli* BW25113 wt whole cell lysate. Lane 2 – Negative control , unload $Ni^{2+}$-agarose resin incubated with *E.coli* whole cell lysate. Lane 3 – His-*Xf*YbbN bound to $Ni^{2+}$-agarose resin incubated with *E.coli* whole cell lysate . Lane 4 – His-*Ec*YbbN bound to $Ni^{2+}$-agarose resin incubated with *E.coli* whole cell lysate. Lane 5 – Negative control, His-PaOhr bound to the $Ni^{2+}$-agarose resin incubated with *E.coli* whole cell lysate.

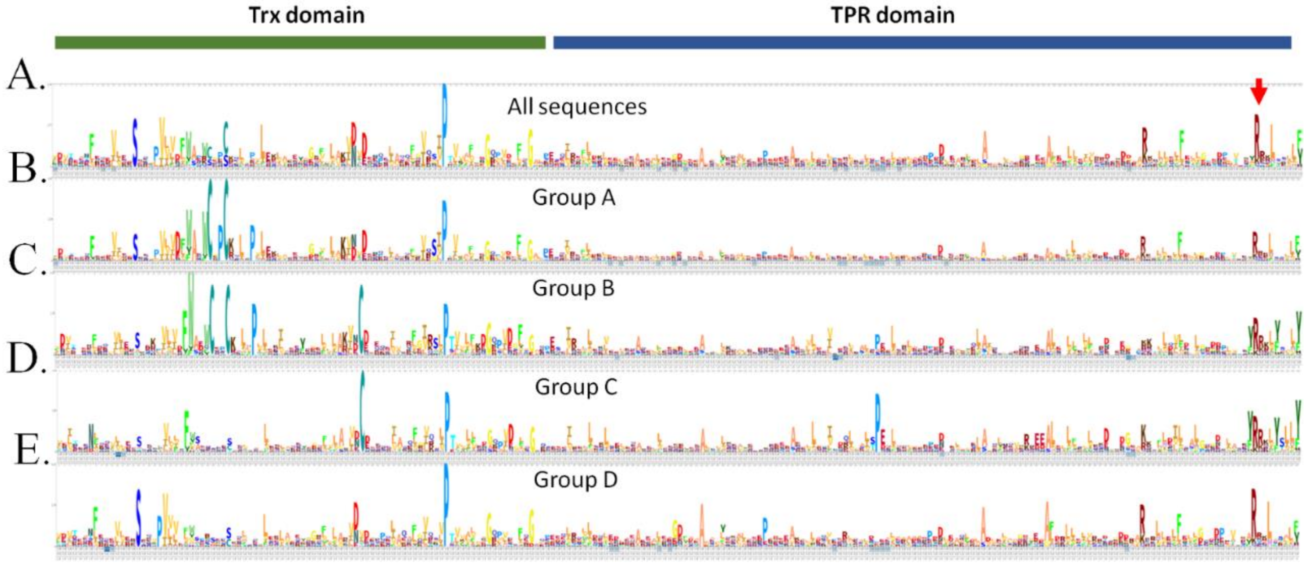


Supplementary Fig. 4. **Graphical representation of both thioredoxin and TPR domains from *Xf*YbbN/CnoX homologue sequences.** Multiple sequence alignment was used to produce a profile hidden Markov model (profile HMM). All profile HMM were visualized by HMM logo created with Skylign. **(A)** HMM logo of all 5532 representative sequences (95 % identity cut-off) extracted from UniProtKB database. **(B)** HMM logo of 3768 sequences representative of *Xf*YbbN group A (CXXC[$N_{24}$]X). **(C)** HMM logo of 219 sequences representative of *Pa*YbbN group B (CXXC[$N_{24}$]C). **(D)** HMM logo of 537 sequences representative of *Ec*YbbN group C (SXXC[$N_{24}$]C). **(E)** HMM logo of 1009 sequences representative of a group D that have CXXC motif is degenerated and lacks the additional Cys residue.

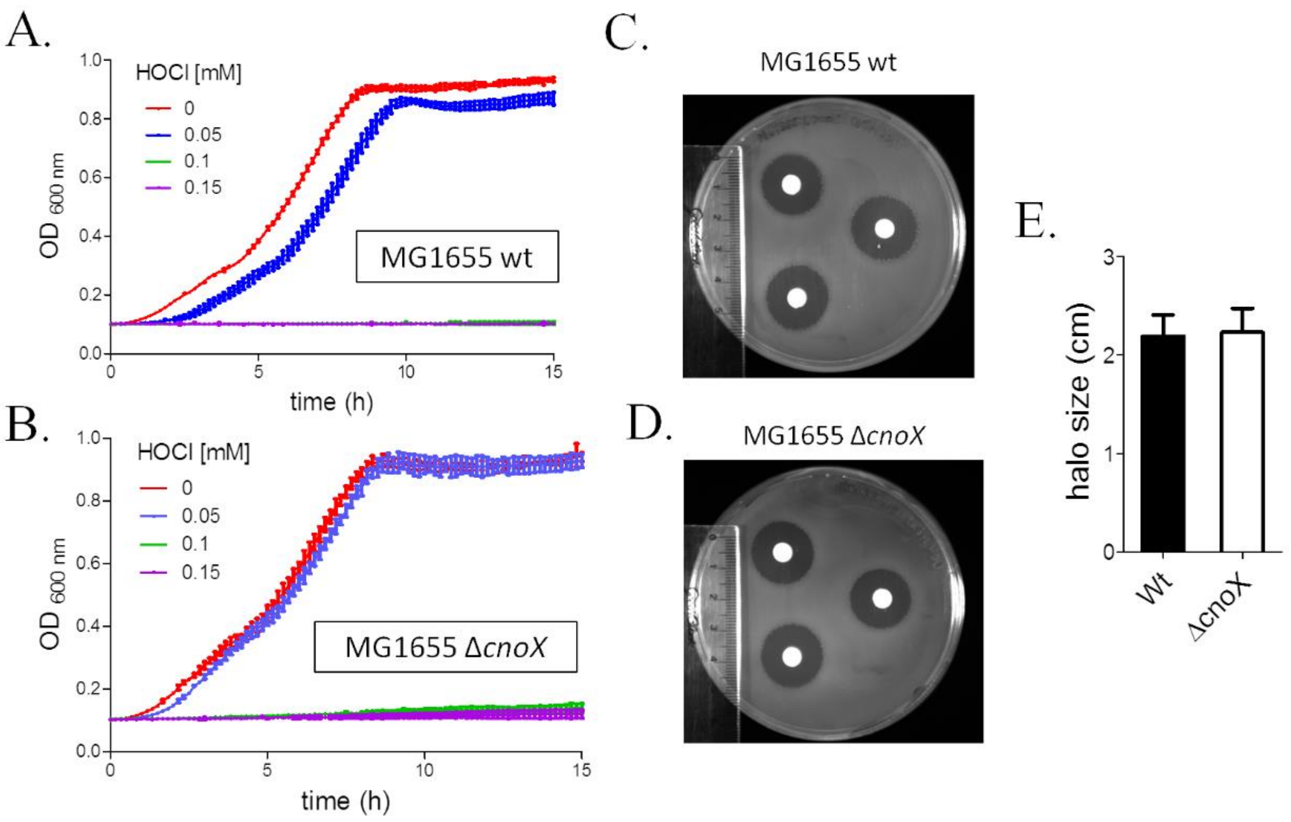


Supplementary Fig. 5. **Effect of HOCl treatment on MG1655 wild type and Δ*ybbN* strains.** Growth curves of MG1655 wild type **(A)** and Δ*ybbN* strains **(B)** in the presence of increasing concentrations of HOCl. To perform the assay, saturated cultures of wild type and Δ*ybbN* strains were diluted in M9 medium supplemented with 0.2 % glucose to $OD_{600}$ 0.1. After that, 180 µL of diluted cultures were distributed in separated wells of 96 well plates and incubated with increasing concentrations of HOCl. The final concentration of HOCl used are indicated in the graphs. Plates were incubated at 37 °C under constant agitation (282 rpm), and the growth profile was monitored during 15 h. Each growth curve represent the media of three technical replicates from three independent biological replicates**.** Kirby-Bauer assay of MG1655 wild type **(C)** and Δ*ybbN* strains **(D)**. The strains were grown until mid-log phase and 100 µL of cell suspension plated on LB and let dry in the laminar flow hood. After that, 6 mm paper filter disks were saturated with 10 µL of solution of 0.5 M of HOCl and placed on the surface of inoculated LB plates. **(E)** Statistical analysis ($n$=12) showed that sensitivity to HOCl wt and *ybbN* mutant strains were not significantly different ($P < 0.01$).

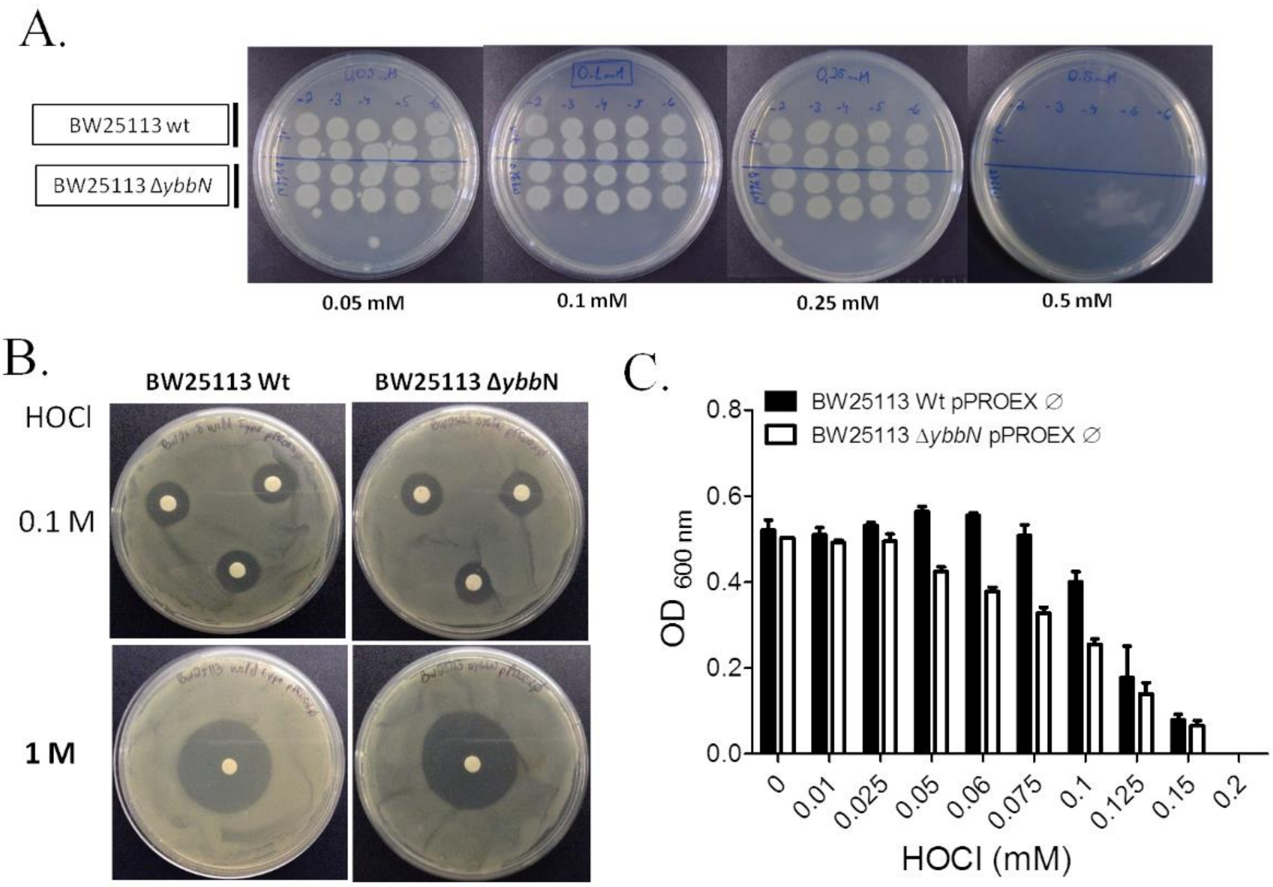


Supplementary Fig. 6. **Effect of HOCl treatment in BW25113 wild type and Δ*ybbN* strains. A.** The cultures were grown in M9 + glucose medium at 37 °C until reach $OD_{600nm}$ 0.2-0.3 and then, treated with different HOCl concentrations for 10 minutes. After, 10 µL of serial dilutions ($10^{-2}$, $10^{-3}$, $10^{-4}$, $10^{-5}$ and $10^{-6}$) were spotted in LB plate and incubated overnight at 37 °C. **B.** Kirby-Bauer assay. The strains were grown until $OD_{600nm}$ 1, plated on LB and let to dry in the laminar flow hood . After that, 6 mm paper filter disks were saturated with 10 µL of solution of either 0.1 or 1 M of HOCl and placed on the surface of inoculated LB plates. **C.** Saturated cultures of wild type and Δ*ybbN* strains harboring empty pPROEX vector were diluted to $OD_{600nm}$ 0.05 in M9 + glucose (0.2%) + ampicillin (100 µg/mL). After that, 180 µL of diluted cultures were distributed in separated wells of 96 well plates and incubated with increasing concentrations of HOCl. The final concentration of HOCl used is indicated in the graph. Plates were incubated at 37 °C under constant agitation and the end-point $OD_{600}$ measured after 16 hour of incubation. The graph is representative of three independent biological replicates**.**

**Supplementary Table 1. Strains, plasmids and oligonucleotides used in this work**

| Strains | Characteristics | Reference |
|---|---|---|
| ***P. aeruginosa*** | | |
| PA14 | Clinical isolate UCBPP-PA14 | [1] |
| *ybbN* mutant | PA14 Δ*ybbN* (*in frame*) | This study |
| ***E. coli*** | | |
| BW25113 | F$^-$ LAM$^-$ *rrn*B3 DE*lac*Z4787 *hsd*R514 DE(*ara*BAD)567 DE(*rha*BAD)568 *rph*-1 | [2] |
| BW25113 Δ*ybbN* | F$^-$ LAM$^-$ *rrn*B3 DE*lac*Z4787 *hsd*R514 DE(*ara*BAD)567 DE(*rha*BAD)568 *rph*-1 *ybbN* Kn$^S$ | [3] and this study |
| K-12 MG1655 | *F$^-$, lambda$^-$, rph-1* | [4] |
| K-12 MG1655 Δ*cnoX* | *F-, lambda-, rph-1 cnoX* | [4] |
| DH5α | *supE44 lacU169*(80 *lac*ZM15) *hsdR17recA1 endA11 gyrA96 thi-1 relA1* | Invitrogen |
| BL21 (DE3) | F$^-$*ompTgal dcm lon* hsd $S_B(r_B^- m_B^-)$ λ(DE3 [*lac*I*lac*UV5-T7 gene 1 *ind*1 *sam*7 *nin*5]) | [5] |
| S17-1 | *F-, RP4-2(Km::Tn7,Tc::Mu-1), pro-82, LAMpir, recA1, endA1, thiE1, hsdR17, creC510, pLFX* | [6] |
| **Plasmids** | | |
| pUCp18 | Stable expression vector used in *Pseudomonas spp.*, Ap$^r$ | [7] |
| pEX18Ap | Suicide vector used for gene replacement in *Pseudomonas spp.*, Ap$^r$ | [8] |
| pET28a-*XfybbN* | Expression vector, Kn$^r$ | PROMEGA |
| **Oligonucleotides** | | |
| PaYbbN_L_NdeI | AGCTCATATGAGCGATACGCCCTACA | This study |

| | | |
|---|---|---|
| PaYbbN_R_BamHI | AGCTGGATCCTCAGTAGAGCGCCTGGTACAG | This study |
| XfybbN_L_NdeI | CGCCATATGTCCGATAAATCC | This study |
| XfybbN_R_XhoI | CGCCTCGAGTCAAAACAAC | This study |
| MutC40A_ For | CTTGGTGTGCTCCGGCCCGCTCGCTGATTC | This study |
| MutC40A_Rev | GAATCAGCGAGCGGGCCGGAGCACACCAAG | This study |
| MutC37AFor | GATTTTTGGGCGCCTTGGAGCGCTCCGTGCCGCTCGCTG | This study |
| MutC37ARev | CAGCGAGCGGCACGGAGCGCTCCAAGGCGCCCAAAAATC | This study |
| MutS35C_For | CTATTTTTGGTCTGAACGTCAGCACTGTTTGCAGTTAAC | This study |
| MutS35C_Rev | GTTAACTGCAAACAGTGCTGACGTTCAGACCAAAAATAG | This study |
| del_YbbN_L_HindIII | GATCAAGCTTAGATGGCACTGCTGGTGAT | This study |
| del_YbbN_R_NdeI | GATCCATATGGTAGGGCGTATCGCTCATGT | This study |
| del_YbbN_L_NdeI | GATCCATATGGCGCTCTACTGAGTCGCC | This study |
| del_YbbN_R_EcoRI | GATCGAATTCTCGCAACTGAGCTGGTAATG | This study |
